# Dynamic Aha1 Co-Chaperone Binding to Human Hsp90

**DOI:** 10.1101/550228

**Authors:** Javier Oroz, Laura J. Blair, Markus Zweckstetter

## Abstract

Hsp90 is an essential chaperone that requires large allosteric changes to determine its ATPase activity and client binding. Because of the inherent low ATPase activity of human Hsp90, the co-chaperone Aha1, which is the only known ATPase stimulator in eukaryotes, is important for regulation of Hsp90’s allosteric timing. Little is known, however, about the structure of the Hsp90/Aha1 full-length complex. Here, we characterize the solution structure of unmodified human Hsp90 in complex with Aha1 using NMR spectroscopy. We show that the 214 kDa complex adopts multiple conformations in the absence of nucleotide. Interaction with Aha1 induces structural changes near the nucleotide-binding site in Hsp90’s N-terminal domain, providing a basis for its ATPase-enhancing activity. Moreover, the E67K mutation in Aha1 strongly diminishes the interaction, supporting a two-step binding mechanism. Our data reveal important aspects of this pivotal chaperone/co-chaperone interaction and emphasize the relevance of characterizing dynamic chaperone structures in solution.

## Introduction

Heat Shock Protein of 90 kDa (Hsp90) is a highly conserved ATP-dependent molecular chaperone responsible for the stabilization, maturation and activation of many client proteins [1,2]. Several co-chaperones regulate Hsp90’s activation cycle, which is essential to maintain protein homeostasis [3,4]. During its activation cycle, the Hsp90 dimer undergoes large conformational rearrangements from an extended to a closed, ATPase-active conformation, a process which involves intra- and inter-protomer interactions [5–7]. Different Hsp90 orthologs show distinct conformational equilibria[8] that are translated into different ATPase activities. Human Hsp90 predominantly populates extended, ATPase-incompetent dimeric conformations [7,8] and, compared to other orthologs, contains a very low inherent ATPase activity [9], despite the crucial role of Hsp90 ATPase activity for cell viability [10].

Aha1 (*Activator of Hsp90 ATPase activity* 1) [11] binds with an affinity of 0.5 mM to human Hsp90 [12]. The Aha1 co-chaperone is unique in its ability to strongly enhance the inherently low ATPase activity of human Hsp90 and, thus, plays an important role as the facilitator for human Hsp90 to fulfill its activation cycle [13]. Specifically, the Hsp90 dimer must close and both N-terminal ATPase domains (Hsp90N) reposition to a closed state for ATP hydrolysis [4]. Several intermediate steps in the closure process of Hsp90 were determined [4]. Aha1 specifically helps to overcome the rate-limiting structural changes in Hsp90, which leads to a potent stimulation of its ATPase activity [4,9,13]. However, the structural basis for this stimulation remains unknown.

Hsp90 is a very dynamic molecule [5], and its N-terminal ATPase domain can freely rotate in solution [14]. Aha1 was suggested to bind asymmetrically to the Hsp90 dimer and thus stabilize the interaction between the two Hsp90 N-terminal domains in the closed dimer structure [15], in a nucleotide-independent manner [16]. Such a static model, however, is difficult to reconcile with the *cis* (Aha1 is bound to one protomer of the Hsp90 dimer) and *trans* (Aha1 is bound to both protomers of the dimer) interactions observed between Aha1 and Hsp90 [15], especially considering the high degree of rotational freedom within Hsp90’s N-terminal domains [14]. In addition, Aha1 was reported to promote a partially and not fully closed Hsp90 dimer conformation [13,15]. Only in presence of nucleotide, Aha1 stabilizes the N-terminally dimerized state of Hsp90 impeding the rotation of these domains [14]. These observations suggest that the interaction between Hsp90 and Aha1 is complex and probably involves multiple transition states promoted by Aha1- and nucleotide-binding, as well as ATP hydrolysis [16,17].

## Results and Discussion

To gain insight into the structural basis of the allosteric binding of Aha1 to Hsp90, we studied the 214 kDa Hsp90/Aha1-complex by solution NMR spectroscopy using methyl-labeled human Hsp90β. Relaxation-optimized NMR spectroscopy in combination with selective labeling has previously been used successfully to characterize dynamic, multi-step interactions between Hsp90 and co-chaperones and/or clients [7,18]. Addition of Aha1 to full-length Hsp90 induced strong changes in the Hsp90 isoleucine NMR spectra (Figure 1a-b), suggesting conformational rearrangements in Hsp90 dimer upon Aha1-induced closure [7,18]. Aha1-specific changes included chemical shift perturbations and cross-peak broadening, but also the appearance of new Hsp90 cross-peaks (Figures 1b, EV1).

**Figure 1.**
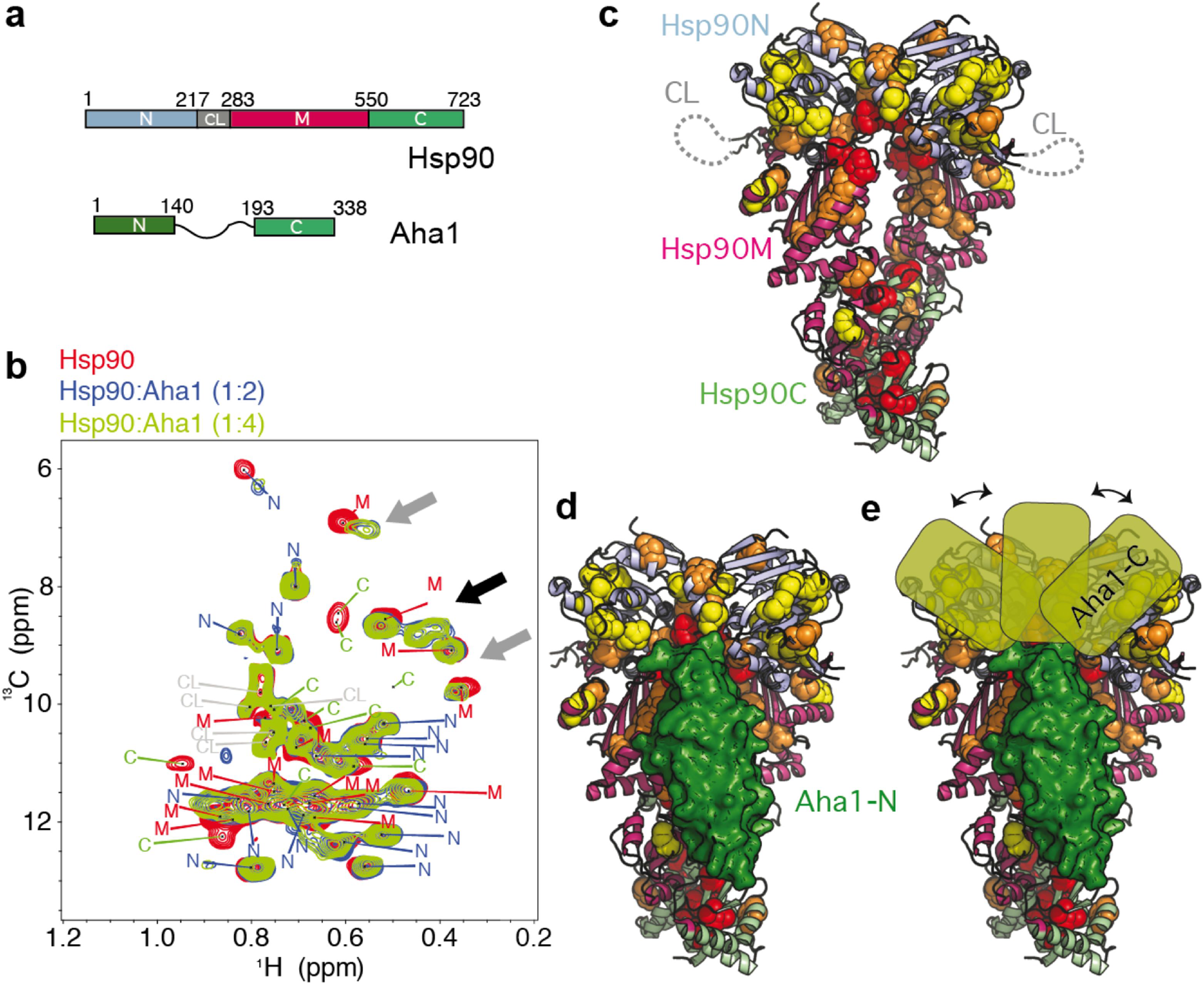
Interaction of Aha1 with human Hsp90 in solution. **a)** Domain organization of human Hsp90β and Aha1. The N-terminal domain (Hsp90N) contains the ATPase active site, Hsp90C is responsible for dimerization. The domain colour code is kept in all figures. **b)** Aha1 binding promotes strong changes in the methyl-TROSY NMR spectra of Hsp90, including chemical shift perturbations and line broadening of Hsp90 isoleucine moieties (grey arrows) and the appearance of new signals (black arrow). The assignment of cross-peaks to specific Hsp90 domains (Hsp90N, Hsp90M and Hsp90C) is indicated. **c)** Distribution of affected Hsp90 isoleucine residues in Aha1 interaction and allostery on the structure of the closed Hsp90 dimer (PDB id 5fwk) [19]. Red, orange and yellow spheres represent NMR perturbations of decreasing magnitude (shown in Figure EV1b). The flexible charged linker of Hsp90 is represented by a dotted line. **d)** Location of Aha1-N (in green surface representation) on the structure of Hsp90 (PDB id 1usu [21]). **e)** The Aha1-affected regions of Hsp90N suggest multiple bound conformations of the C-terminal domain of Aha1 (Aha1-C).

Mapping of the affected isoleucines on the structure of Hsp90 [19] showed that the most strongly affected residues are located at the interface between the middle (Hsp90M) and C-terminal domain (Hsp90C) of Hsp90, as well as the N-terminal interface involved in dimer closure (Figure 1c). Both interfaces undergo rearrangements during Hsp90 allosteric changes [20]. In addition, the regions affected in Hsp90M agree with the binding site of the N-terminal domain of Aha1 (Aha1-N), as displayed in the crystal structure of the complex of yeast Aha1-N with Hsp90M (Figure 1c-d) [21]. Hsp90N, however, which would interact with Aha1-C assuming a head-to-tail mode of interaction [21], was more broadly affected (Figures 1c-d, EV2), in agreement with the footprinting of the Hsp90/Aha1-interaction using crosslinking coupled to mass spectrometry [12], but contrary to a previously proposed model [15]. Because one Aha1 molecule binds to one Hsp90 dimer [15], the broad distribution of chemical shift perturbations in Hsp90N suggests that Aha1-C binds in *cis* and *trans* [15] to multiple sites of both Hsp90N domains of the Hsp90 dimer leading to different Hsp90/Aha1-complex structures (Figure 1e). The dynamic nature of the interaction may be supported by the structural flexibility that Hsp90N retains in complex with Aha1 in the absence of nucleotide [14] and could further be favored by the flexibility of the long interdomain linker in Aha1 (Figure 1a). Notably, the interdomain linker in Aha1 is positively charged (pI = 8.3), while the flexible linker (CL) [22] between the N- and M-domains of Hsp90 is negatively charged (Figure 1a, c; pI = 4.5), favoring electrostatic interactions between these parts of Aha1 and Hsp90. In addition to the polymorphic interactions on Hsp90N (Figure 1), Aha1 binding can induce conformational rearrangements on Hsp90N involving distant regions of the domain (Figures EV2, EV3).

To discriminate between the binding of Aha1 and subsequent allosteric rearrangements induced in Hsp90 by the co-chaperone, we prepared an Hsp90 construct containing only the N-terminal and middle domains of Hsp90 (Hsp90NM), i.e. the Hsp90 regions that are most relevant for the interaction (Figure 1e). Because Hsp90NM is monomeric in solution [7], it cannot undergo the allosteric changes involved in dimer closure. Aha1 will therefore only be able to form *cis* interactions with Hsp90NM (Figure 2). NMR data of the interaction show that Aha1-binding was strongly attenuated when compared to full-length Hsp90 (Figure EV4a-c). Small-angle X-ray scattering (SAXS) of an equimolar Hsp90NM/Aha1-mixture also resembled the scattering profile expected for a mixture of isolated proteins rather than for a stable complex (Figure EV4f). The Hsp90NM/Aha1-interaction is thus weak, in agreement with a previous report suggesting that Aha1/Hsp90 *trans* interaction is more stable than Aha1/Hsp90 *cis* interaction [12]. The combined data show that Hsp90/Aha1 *trans* interactions are important for complex stabilization [12], although Aha1 bound in *cis* to the full-length Hsp90 dimer might reach the same level of ATPase stimulation [15].

**Figure 2.**
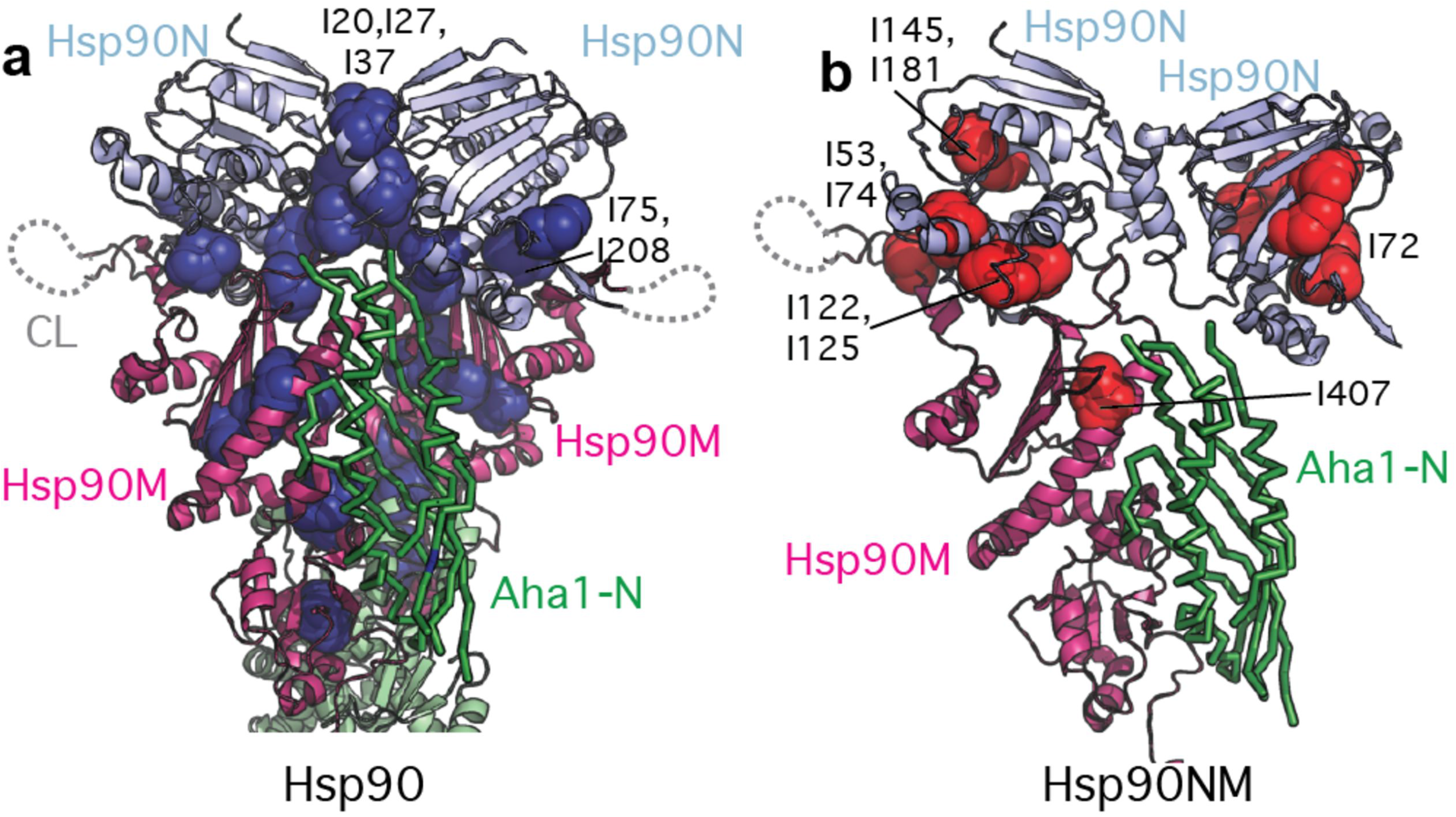
Discrimination between allosteric changes (a) and *cis* binding (b). **a)** Mapping of residues that are more affected in the Hsp90/Aha1-complex compared to the Hsp90NM/Aha1-interaction (dark blue spheres, blue bars in Figure EV4d) on the structure of the closed Hsp90 dimer. **b)** Residues that are more affected in the Hsp90NM/Aha1-interaction when compared to the Hsp90/Aha1-complex (red bars in Figure EV4d) are displayed with red spheres on Hsp90NM. To allow better comparison with panel (a), a second Hsp90N domain is displayed (as observed in the closed Hsp90 dimer; Figure 1c), as well as the N-terminal domain of Aha1 (green).

Comparison of the perturbations induced by Aha1 in Hsp90 with those induced in Hsp90NM (Figure EV4d) provide the unique opportunity to distinguish between Hsp90 regions affected only in the *cis* binding process (Figure 2b) and those experiencing allosteric changes in the complex upon dimer closure (Figure 2a). Aha1 binding induces stronger effects on Hsp90M moieties for full-length Hsp90 (Figure EV4d), supporting the weaker binding of Aha1 to Hsp90NM. Remarkably Hsp90N shows two clear interfaces: residues I20, I27, I37, I75 and I208 are more affected in the Hsp90/Aha1-complex (Figures 2a, EV4d). These residues are located in the Hsp90N dimer interface (I20, I27 and I37), and in the Hsp90N/M interface (I75 and I208), and thus reflect allosteric rearrangements. On the other hand, I53, I72, I74, I122, I125, I145 and I181 were more strongly broadened in the Hsp90NM/Aha1-interaction and form a continuous region in Hsp90N opposite to the dimer interface (Figures 2b, EV4d), supporting the polymorphic interaction between the C-terminal domain of Aha1 and the N-terminal domain of Hsp90 (Figure 1e). Because Hsp90N can freely rotate even in complex with Aha1 [14], the *cis* binding interface shown in Figure 2b could be rotated 180° upon Aha1-binding.

To confirm the importance of the interaction of the N-terminal domain of Aha1 with the middle domain of Hsp90, as seen in the complex structure of the two isolated domains [21], we characterized the binding of the mutant protein E67K-Aha1 to Hsp90. E67K-Aha1 is unable to stimulate the ATPase activity of Hsp90 due to impaired binding [12] and is ineffective in promoting Hsp90-dependent maturation [12] or aggregation of clients [23]. Consistent with these data, binding of E67K-Aha1 to human Hsp90 was weaker and did not result in new Hsp90 cross-peaks (Figures EV5a-b), in contrast to wild-type Aha1 (Figure 1b). This effect can be rationalized based on the atomic structure of the yeast Hsp90M/Aha1-N-complex [21]: Residue E67 of Aha1 is surrounded in the complex by a cluster of positively charged residues (K398, K400, R404 and K405) of Hsp90M (Figure EV5e) such that the E67K mutation results in electrostatic repulsion.

Binding of wild-type Aha1 to full-length Hsp90, but not to Hsp90NM, resulted in new Hsp90 cross-peaks in the absence of nucleotide (Figure EV4). The new Hsp90 signals could be due to inter-protomer contacts established between isoleucines in the Hsp90N dimerization interface upon Aha1-induced dimeric closure of Hsp90 (Figure 1c), or due to intra-protomer conformational rearrangements in the chaperone induced by co-chaperone action (Figure EV3). To distinguish between these possibilities, we compared the Hsp90 spectrum in presence of Aha1 (without nucleotide) with the Hsp90 spectrum when both the ATPase-inhibiting co-chaperone FKBP51 [7] and nucleotide were present (Figure 3a-c). In the latter conditions, the Hsp90 dimer will be extended, because FKBP51 impedes the allosteric closure of Hsp90 (Figure 3e-f) [7]. The comparison showed that the new set of cross-peaks, which appear upon Aha1-binding in the absence of nucleotide (Figure 3a), partially resemble the new set of cross-peaks that appear in the Hsp90/ADP/FKBP51 (Figure 3b) and Hsp90/AMP.PNP/FKBP51 complexes (Figure 3c). In agreement with previous results [4], SAXS analysis of the Hsp90/Aha1-complex showed that co-chaperone binding induced a partially closed conformation of Hsp90 in both the absence and presence of AMP.PNP (Figure 3d). The comparison suggests that the Aha1-driven, complex-specific Hsp90 signals arise from conformational changes induced in Hsp90 by Aha1-binding, which are independent of inter-protomer allosteric closure.

**Figure 3.**
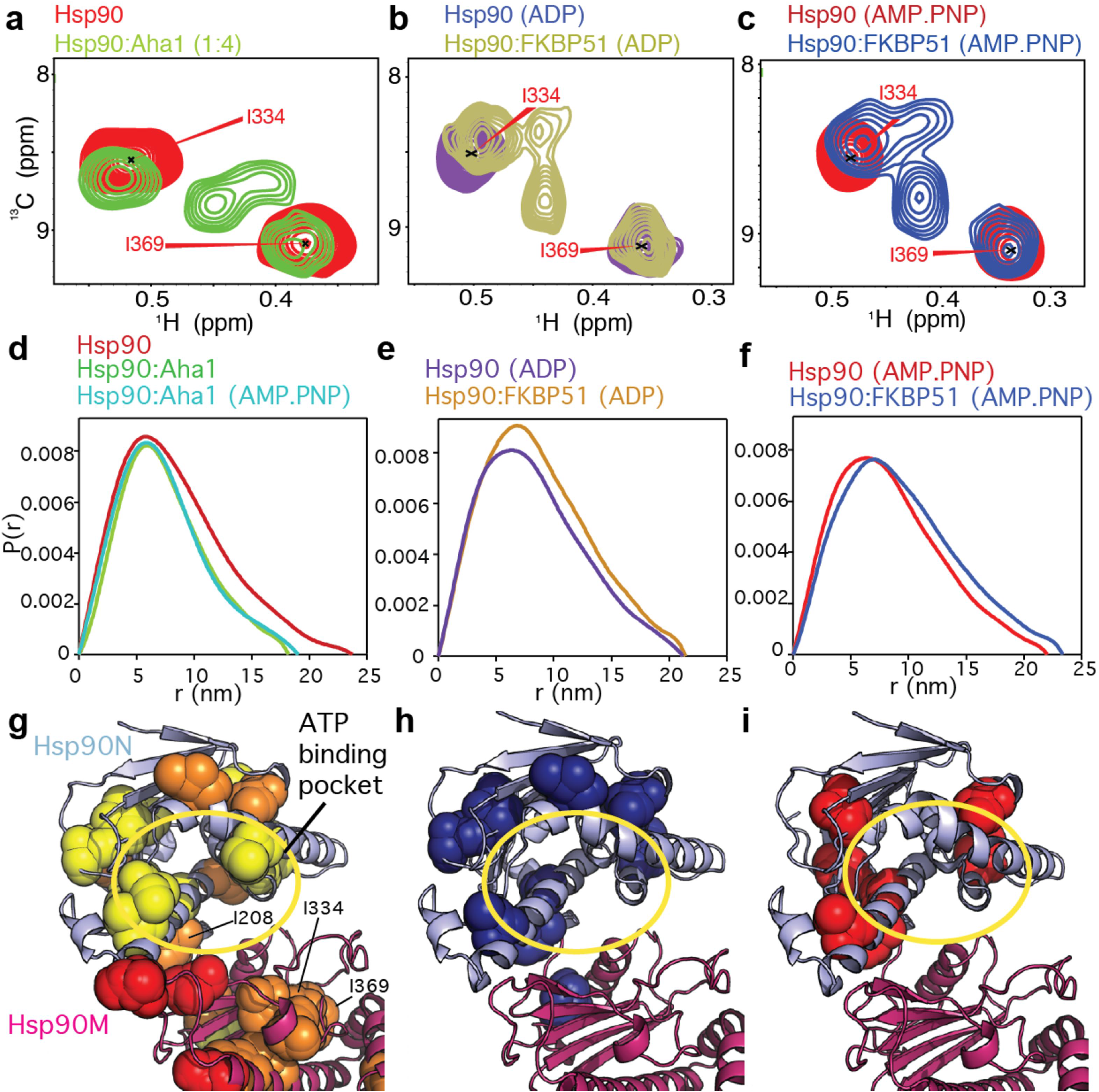
Structural changes induced by Aha1 and nucleotide binding. **a)** New signals appear upon Aha1-binding in the methyl-TROSY spectra of full-length Hsp90. **b-c)** New cross peaks appear upon binding of nucleotide and FKBP51. These signals are similar for ADP and AMP.PNP, but are partially different from the peaks appearing in the presence of Aha1 (a). **d)** SAXS P(r) distribution shows that Aha1 promotes a partially closed conformation [13,15] of Hsp90 independent of nucleotide. **e-f)** FKBP51 stabilizes the extended conformation of Hsp90 [7]. **g)** Aha1 promotes structural changes around the ATP-binding pocket in Hsp90N in the absence of nucleotide. **h-i)** Nucleotide binding promotes structural rearrangements around the ATP-binding pocket [7]. Spheres in **(g-i)** represent isoleucine residues affected in the corresponding binding process. A yellow circle in **(g-i)** highlights the ATP-binding pocket.

Binding of nucleotides to Hsp90 induces a rotation of Hsp90N [14,24] and structural changes in the ATP lid [5,20] (Figure EV3a), which promote the trapping of the nucleotide in the binding pocket [25]. The spectral changes of Hsp90 upon Aha1 binding in the absence of nucleotide are similar, but not identical to those induced by nucleotide binding (Figure 3a-c, EV3). This suggests that Aha1 induces intermediate conformational rearrangements around the ATP-binding pocket of Hsp90N, which still allow for ATP exchange [13] but that energetically facilitate the additional changes required for ATP trapping and hydrolysis [26,27]. Consistent with this hypothesis, NMR spectroscopy indicated that Aha1 promotes rearrangements on Hsp90N that resemble those induced by Hsp90N dimerization (Figure EV3b), preparing the domain for nucleotide trapping.

Our data support the multi-step nature of the Hsp90/Aha1-interaction [17] and indicate that the C-terminal domain of Aha1 can adopt several conformations leading to a dynamic, polymorphic complex. We further showed that Aha1 promotes a partially closed conformation of Hsp90 where the still flexible [14] N-terminal domains of Hsp90 undergo conformational rearrangements towards an intermediate state that facilitates ATP binding. Because Aha1 is critical for accelerating the ATPase activity of human Hsp90 [13], our study helps in deciphering the molecular mechanism of this stimulation and, thus, in the understanding of the activation cycle of Hsp90, which is fundamental to maintain eukaryotic homeostasis [3,4].

## Materials and Methods

Human Hsp90β and Aha1 were cloned into pET28b and pET21a vectors, respectively (Novagen), and expressed in BL21(DE3) *E. coli* strain. Metabolic precursors for selective [^1^H-^13^C]-labeling of Hsp90 isoleucine d1 methyl groups in fully deuterated media were purchased from NMR-Bio. Selective labelling was achieved as described in [28]. Cells were lysed by sonication and recombinant proteins purified by Ni^2+^ affinity chromatography in Ni-NTA agarose (Thermo Fisher) using 50 mM TrisHCl/500 mM NaCl/10 mM imidazole [ph 8.0] as binding buffer, increasing to 250 mM imidazole for elution. Aha1 was subjected to tobacco etch virus (TEV) proteolysis and subsequently purified again by Ni^2^+ affinity purification. Proteins were further purified by size exclusion chromatography in 10 mM Hepes/500 mM KCl/1 mM DTT [pH 7.5] using a Superdex 200 column (gE Healthcare).

NMR experiments were acquired at 25 °C on Bruker Avance III 800 MHz and 900 MHz spectrometers (both equipped with TCI cryoprobes) using 50 mM sodium phosphate/300 mM NaCl/1 mM DTT [pH 7.2] in 100% D_2_O. Spectra in the presence of nucleotide were obtained in 20 mM Hepes/5 mM KCl/10 mM MgCl_2_/1 mM DTT [pH 7.4] in 100% D_2_O. 80-100 mM isoleucine-labeled Hsp90 was used for NMR titrations. NMR binding plots show the decay in signal intensity (Figures EV1, EV4–EV5), where I_0_ is the intensity of the cross-peaks in the reference spectra (Hsp90 alone). Assignment of Hsp90 isoleucine d1 methyl groups is described in detail in [7]. Spectra were processed using TopSpin (Bruker) and analyzed in Sparky (Goddard & Kneller, UCSF).

SAXS data were collected at 25 °C from pure and monodisperse samples of Hsp90 and Aha1 in 50 mM sodium phosphate/10 mM NaCl/1 mM DTT [pH 6.8] or 20 mM Hepes/5 mM KCl/10 mM MgCl_2_/1 mM DTT [pH 7.4] in the presence of 2 mM AMP.PNP. Sample concentrations ranged from 20-100 mM, and shown P(r) distributions contained 50 mM Hsp90 and equimolar amounts of Aha1. Scattering profiles were analysed using standard procedures using ATSAS[29]. Data collection for the Hsp90/FKBP51/nucleotide complexes is described in [7]. SAXS measurements were performed at DESY (Hamburg, Germany) and Diamond Light Source (Oxford, UK) stations.

## Acknowledgements

We thank Jeremy D. Baker for providing the cDNA clones of human Aha1 and E67K-Aha1, and Bliss Chang for help with the NMR assignment of Hsp90 isoleucine methyl groups. J.O. was supported by a Marie Curie Intra-European fellowship (project number 626526). M.Z. was supported by the European Community’s Seventh Framework Programme (FP7/2007-2013) under Biostruct-X (grant agreement 283570), and the advanced grant “87679-LLPS-NMR” of the European Research Council.

## Author contributions

J.O. performed the experiments and analyzed the data; J.O., L.J.B. and M.Z. conceived the project; J.O. and M.Z. wrote the manuscript.

## Expanded View

**Expanded View Figure 1.**
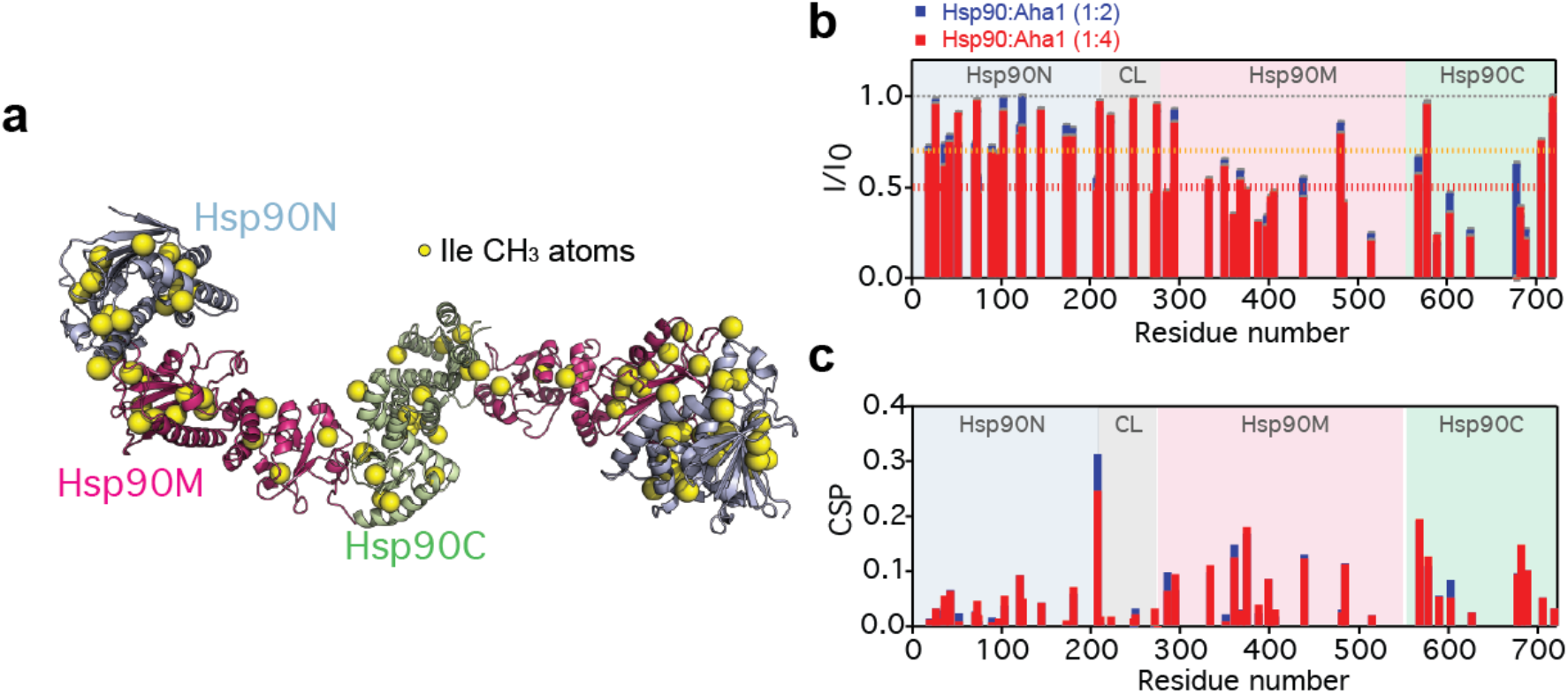
NMR spectroscopy of the Hsp90/Aha1-interaction. **a)** Localization of ^13^C-labeled isoleucine methyl groups (yellow spheres) on the extended conformation of the human Hsp90 dimer [1] (modified from PDB id 5fwk) [2]. **b)** Sequence-specific analysis of the changes in Hsp90 peak intensity upon addition of Aha1. Blue and red bars represent 1:2 and 1:4 molar ratios, respectively. The orange and red lines mark I/I_0_ thresholds of 0.7 and 0.5, respectively, to color code residues in Figure 1c-e: I/I_0_<0.5 in red; I/I_0_= 0.5-0.7 in orange; I/I_0_= 0.7-1 in yellow. Error bars (in grey, very small) were calculated using the spectral signal-to-noise ratios. **c)** Chemical shift perturbation plot of the Hsp90/Aha1-interaction. Hsp90 domains are indicated.

**Expanded View Figure 2.**
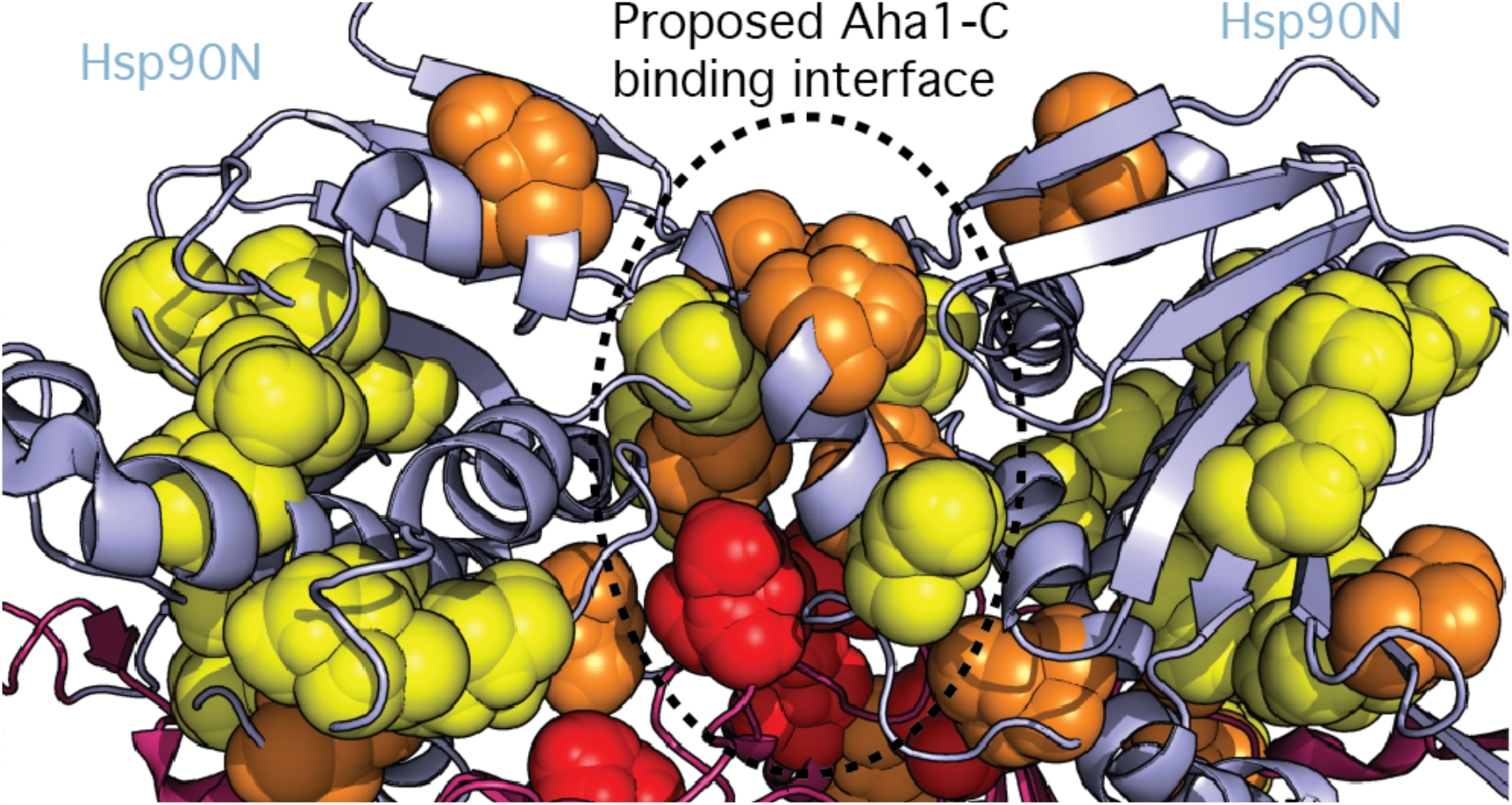
Multiple regions of the N-terminal domain of Hsp90 are affected by Aha1 binding. Isoleucine methyl groups of the N-terminal domain of Hsp90, which are perturbed by addition of Aha1 to full-length Hsp90 (Figure 1b), are represented by spheres using the structure of the closed Hsp90 dimer (PDB id 5fwk). The color code is the same as used in Figures 1 and EV1. The previously proposed Aha1-C binding interface [3] is marked.

**Expanded View Figure 3.**
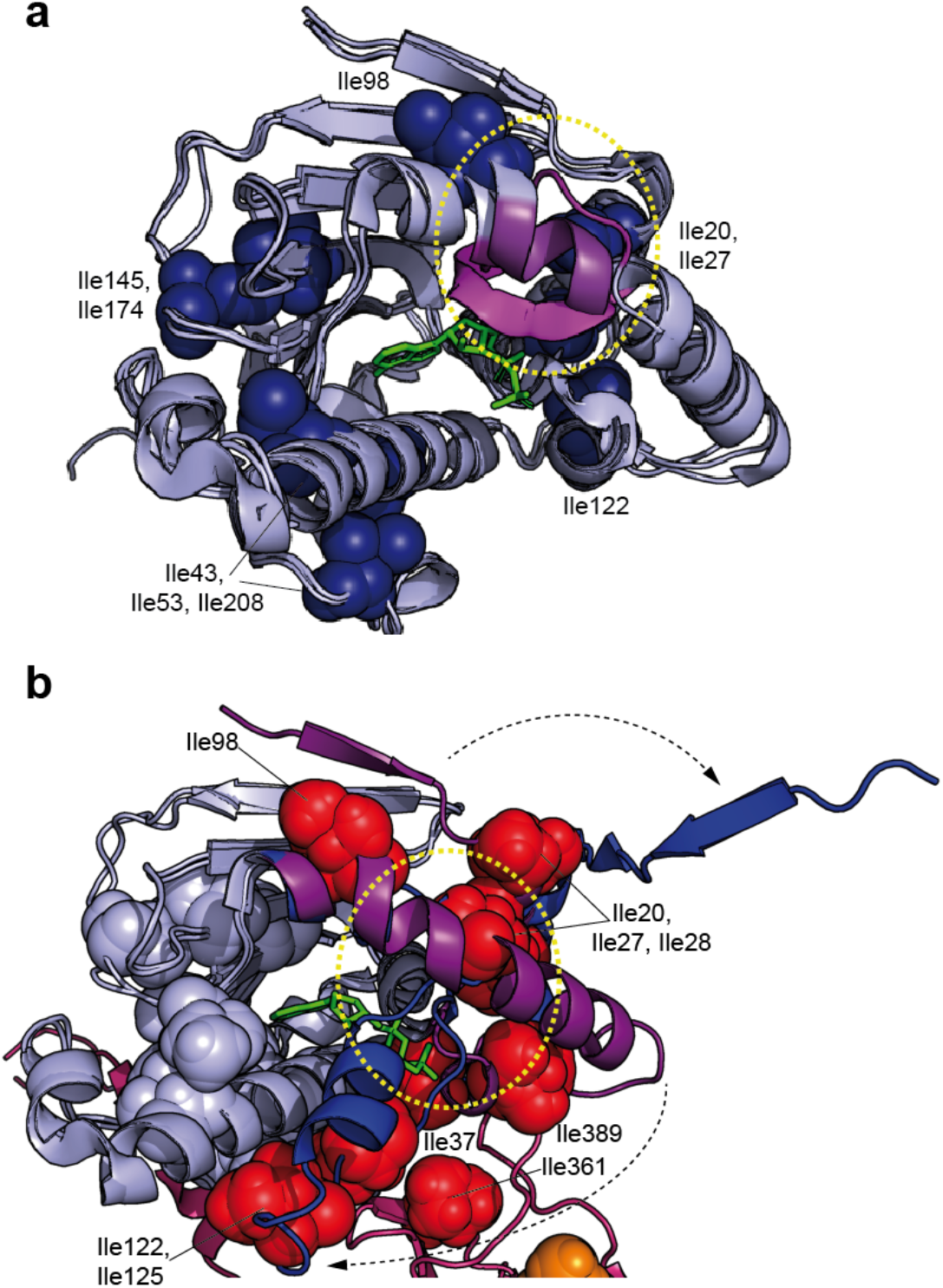
Conformational changes in the N-terminal domain of Hsp90 upon nucleotide binding and dimerization. **a)** Superposition of the structure of the N-terminal domain of Hsp90 (Hsp90N) in the absence of ADP (PDB id: 5j2v) with two structures in the presence of ADP (PDB id: 1am1, ADP-bound yeast Hsp90; 1uym, Hsp90Nb bound to the PU3 analog of ATP). ADP is displayed with green sticks. Nucleotide binding to Hsp90N induces changes in the proximal part of helix 4 (highlighted in magenta, yellow circle). Isoleucine residues, for which the methyl cross-peaks are perturbed by binding of ADP to full-length Hsp90 (please see Figure 3h), are displayed with blue spheres. **b)** Superposition of monomeric Hsp90N structure (PDB id: 1uym) with Hsp90N in the Hsp90 dimer (PDB id: 5fwk), revealing large conformational changes (indicated by arrows) during dimer closure and Hsp90N dimerization: β-Strand 1 swaps between domains and helix 4 folds around the hinge (yellow circle) to fully close the nucleotide binding pocket. Regions undergoing strong conformational changes are marked in magenta and dark blue in monomeric and dimeric Hsp90N, respectively. Isoleucine residues, for which the methyl cross-peaks are perturbed upon addition of Aha1 to full-length Hsp90 in the absence of nucleotide (Figures 1, EV1) are displayed by spheres (red and light blue). I20, I27, I28, I37, I98, I122 and I125 probe conformational rearrangements in Hsp90N upon dimerization, while I361 and I389 are located on the Hsp90M loop that clamps the new conformation of Hsp90N’s helix 4. All these isoleucine residues are affected upon Aha1 binding, suggesting that the co-chaperone induces conformational rearrangements in Hsp90N similar to those induced by nucleotide binding and dimerization. Light blue spheres mark the Aha1 *cis/trans* binding interface of Hsp90N (Figure 2). ATP is displayed with green sticks.

**Expanded View Figure 4.**
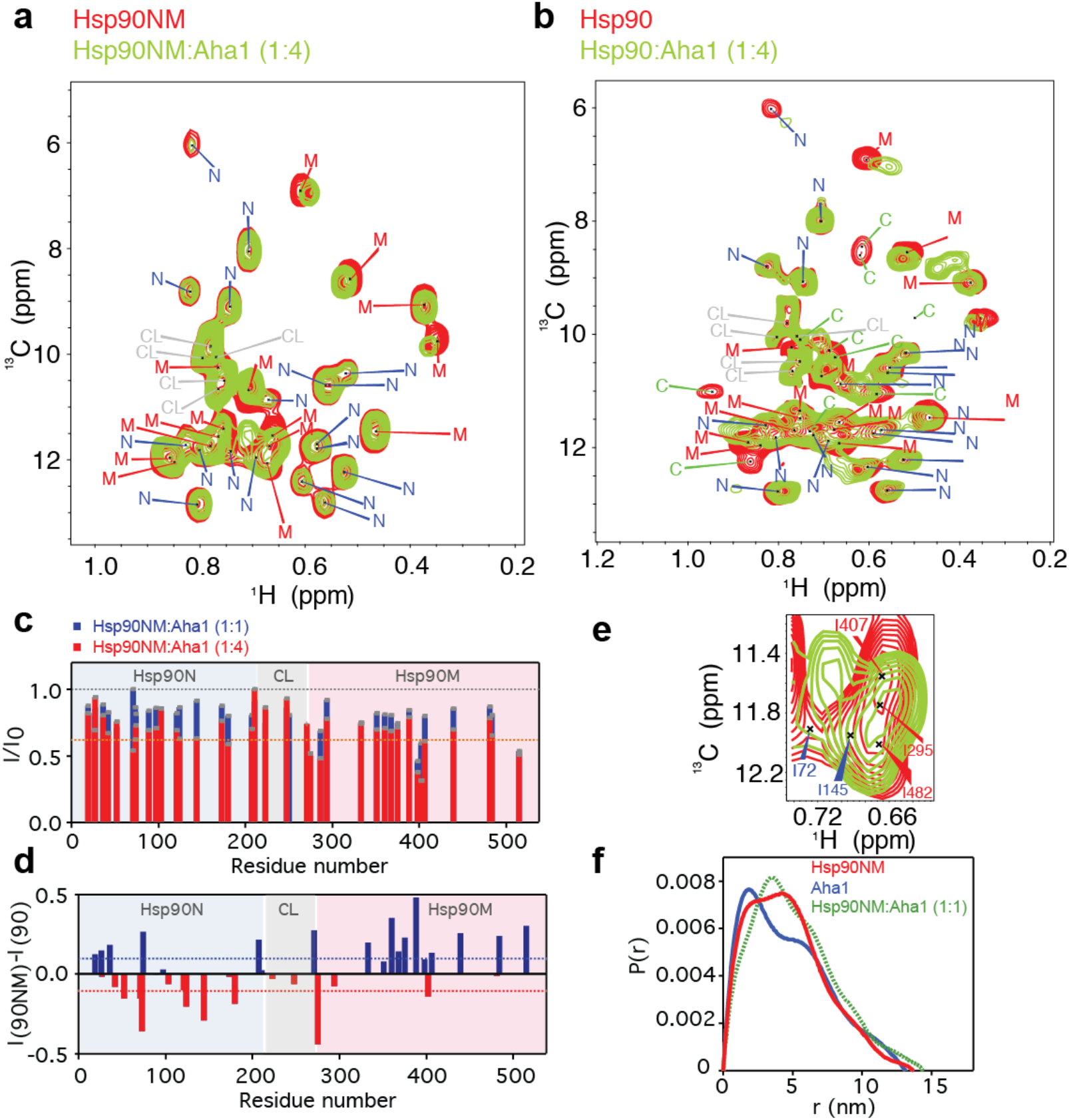
Aha1 binds weakly to monomeric Hsp90NM. **a-b)** Methyl-TROSY spectra of Hsp90NM showing that Aha1-binding does not promote the appearance of new isoleucine cross-peaks (a), in contrast to Aha1-binding to full-length, dimeric Hsp90 (b, same as in Figure 1b). Molar ratios are indicated. **c)** Sequence-specific decrease in the intensities of methyl cross-peaks of Hsp90NM upon addition of equimolar (blue) and 4-fold molar excess (red) of Aha1. An orange dashed line at I/I_0_=0.7 marks the threshold below which Hsp90NM residues are considered to be significantly affected upon addition of Aha1. Error bars (grey bars, very small) are based on spectral S/N ratios. **d)** Comparison of Hsp90/Aha1-with Hsp90NM/Aha1-interaction. (I/I_0_) ratios of Hsp90 isoleucine residues upon Aha1 interaction (Figure EV1b) were subtracted from (I/I_0_) values observed for the Hsp90NM/Aha1-interaction (shown in c; both at a molar ratio of 1:4). Positive values (blue bars) indicate isoleucine residues, which were more strongly broadened in full-length Hsp90 upon addition of Aha1 (Hsp90/Aha1-interaction), while negative values (red bars) indicate residues that were more strongly attenuated in Hsp90NM upon addition of Aha1 (Hsp90NM/Aha1-interaction). Blue and red bars correspond to highlighted residues in (Figure 2). **e)** Selected region of the Hsp90NM methyl-TROSY spectra in the absence (red) and presence (green) of Aha1 (as shown in panel (a)). I72 and I407 were strongly shifted. **f)** SAXS P(r) distribution of the mixture (in green) indicates that the complex is not stable and that it is highly dynamic and/or polymorphic.

**Supplementary Figure 5.**
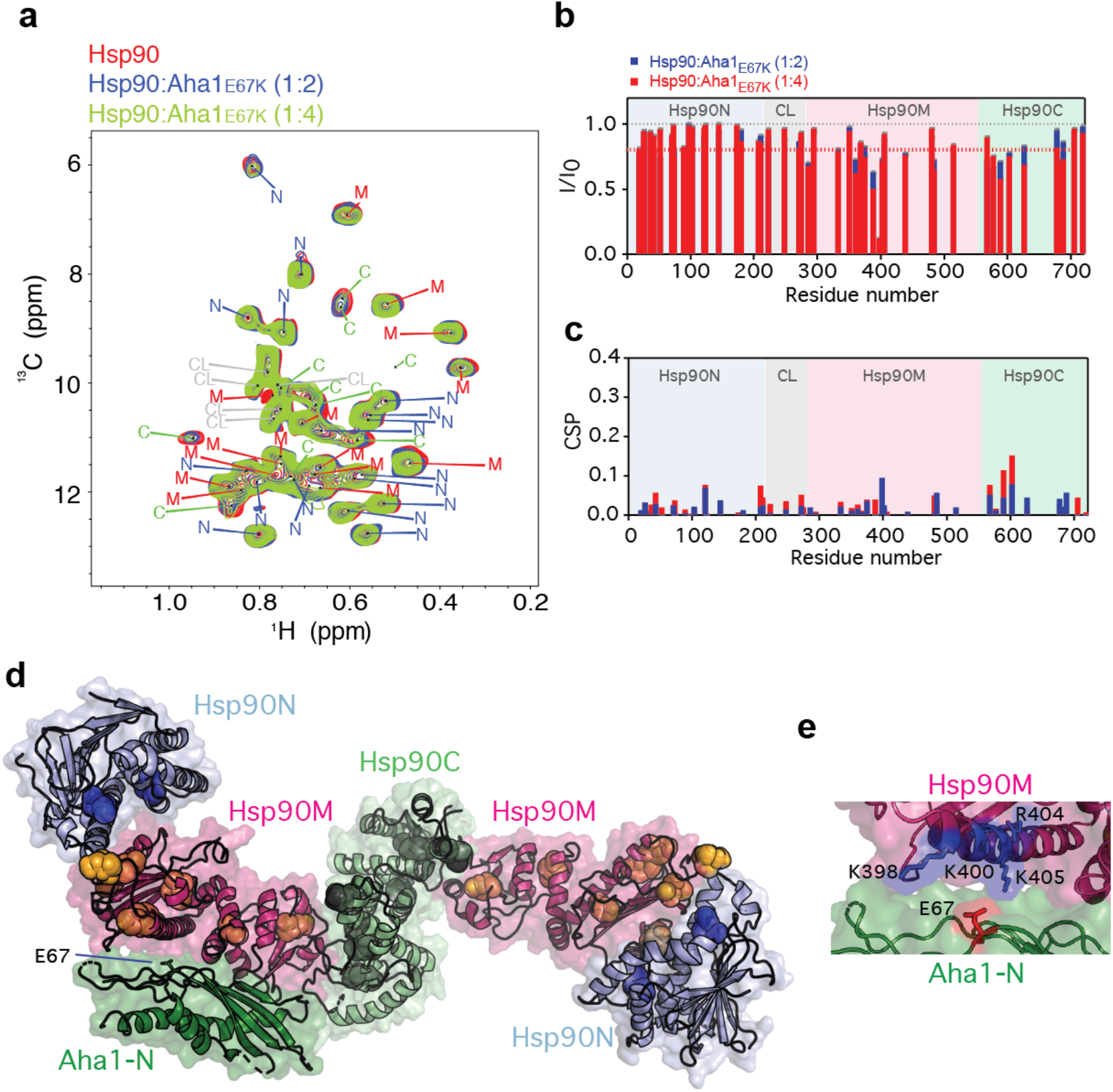
E67K mutation in the N-terminal domain of Aha1 diminishes binding to human Hsp90. **a)** Superposition of isoleucine methyl-TROSY spectra of Hsp90 in the absence (red) and presence of a 2-fold (blue) and 4-fold molar excess of the E67K-mutant Aha1. **b-d)** Sequence-specific changes in the intensity (b) and position (c) of isoleucine methyl cross-peaks of Hsp90 upon addition of the E67K-mutant Aha1 (as displayed in panel (a)). Error bars (in grey, very small) were calculated based on spectral S/N ratios. Isoleucine residues with intensity ratios (I/I_0_) below 0.8 are displayed as spheres in (d), with the N-terminal domain of Aha1 (Aha1-N) positioned according to PDB id 1usu [4]. The Hsp90 dimer is shown in an extended, ATP-incompetent conformation (modified from PDB id 5fwk [2]) following the known inability of the E67K-mutant Aha1 to stimulate Hsp90’s ATPase activity [5]. **e)** The side-chain of E67 (in red sticks) in Aha1-N is surrounded by a cluster of positively charged residues of Hsp90M (K398, K400, R404 and K405, represented in blue sticks).

## References

1. Hartl FU, Bracher A, Hayer-Hartl M (2011) Molecular chaperones in protein folding and proteostasis. Nature 475: 324–332

2. Saibil H (2013) Chaperone machines for protein folding, unfolding and disaggregation. Nat Rev Mol Cell Biol 14: 630–642

3. Taipale M, Jarosz DF, Lindquist S (2010) HSP90 at the hub of protein homeostasis: emerging mechanistic insights. Nat Rev Mol Cell Biol 11: 515–528

4. Schopf FH, Biebl MM, Buchner J (2017) The HSP90 chaperone machinery. Nat Rev Mol Cell Biol 18: 345–360

5. Shiau AK, Harris SF, Southworth DR, Agard DA (2006) Structural Analysis of E. coli hsp90 Reveals Dramatic Nucleotide-Dependent Conformational Rearrangements. Cell 127: 329–340

6. Mayer MP, Le Breton L (2015) Hsp90: Breaking the Symmetry. Mol Cell 58: 8–20

7. Oroz J, Chang BJ, Wysoczanski P, Lee CT, Perez-Lara A, Chakraborty P, Hofele RV, Baker JD, Blair LJ, Biernat J, et al. (2018) Structure and pro-toxic mechanism of the human Hsp90/PPIase/Tau complex. Nat Commun 9: 4532

8. Southworth DR, Agard DA (2008) Species-Dependent Ensembles of Conserved Conformational States Define the Hsp90 Chaperone ATPase Cycle. Mol Cell 32: 631–640

9. Richter K, Soroka J, Skalniak L, Leskovar A, Hessling M, Reinstein J, Buchner J (2008) Conserved conformational changes in the ATPase cycle of human Hsp90. J Biol Chem 283: 17757–17765

10. Panaretou B, Prodromou C, Roe SM, O’Brien R, Ladbury JE, Piper PW, Pearl LH (1998) ATP binding and Hydrolysis are essential to the function of the Hsp90 molecular chaperone in vivo. EMBO J 17: 4829–4836

11. Panaretou B, Siligardi G, P. M, Maloney A, Sullivan JK, Singh S, Millson SH, Clarke PA, Naaby-Hansen S, Stein R, et al. (2002) Activation of the ATPase Activity of Hsp90 by the Stress-Regulated Cochaperone Aha1. Mol Cell 10: 1307–1318

12. Koulov AK, LaPointe P, Lu B, Razvi A, Coppinger J, Dong M-Q, Matteson J, Laister R, Arrowsmith C, Yates III JR, et al. (2010) Biological and structural basis for Aha1 regultaion of Hsp90 ATPase activity in maintaining proteostasis in the human disease cystic fibrosis. Mol Biol Cell 21: 871–884

13. Li J, Richter K, Reinstein J, Buchner J (2013) Integration of the accelerator Aha1 in the Hsp90 co-chaperone cycle. Nat Struct Mol Biol 20: 326–331

14. Daturpalli S, Kniess RA, Lee CT, Mayer MP (2017) Large rotation of the N-terminal domain of Hsp90 is important for interaction with some but not all client proteins. J Mol Biol 429: 1406–1423

15. Retzlaff M, Hagn F, Mitschke L, Hessling M, Gugel F, Kessler H, Richter K, Buchner J (2010) Asymmetric Activation of the Hsp90 Dimer by Its Cochaperone Aha1. Mol Cell 37: 344–354

16. Wortmann P, Götz M, Hugel T (2017) Cooperative nucleotide binding in Hsp90 and the underlying mechanisms. Biophys J 113: 1711–1718

17. Wolmarans A, Lee B, Spyracopoulos L, LaPointe P (2016) The Mechanism of Hsp90 ATPase Stimulation by Aha1. Sci Rep 6: 33179

18. Oroz J, Kim JH, Chang BJ, Zweckstetter M (2017) Mechanistic basis for the recognition of a misfolded protein by the molecular chaperone Hsp90. Nat Struct Mol Biol 24: 407–413

19. Verba K, A., Wang RY-R, Arakawa A, Liu Y, Shirouzu M, Yokoyama S, Agard DA (2016) Atomic structure of Hsp90-Cdc37-Cdk4 reveals that Hsp90 traps and stabilizes an unfolded kinase. Science 352: 1542–1547

20. Krukenberg KA, Street TO, Lavery LA, Agard DA (2011) Conformational dynamics of the molecular chaperone Hsp90. Quat Rev Biophys 44: 229–255

21. Meyer P, Prodromou C, Liao C, Hu B, Roe SM, Vaughan CK, Vlasic I, Panaretou B, Piper PW, Pearl LH (2004) Structural basis for recruitment of the ATPase activator Aha1 to the Hsp90 chaperone machinery. EMBO J 23: 1402–1410

22. Jahn M, Rehn A, Pelz B, Hellenkamp B, Richter K, Rief M, Buchner J, Hugel T (2014) The charged linker of the molecular chaperone Hsp90 modulates domain contacts and biological function. Proc Natl Acad Sci USA 111: 17881–17886

23. Shelton LB, Baker JD, Zheng D, Sullivan LE, Solanki PK, Webster JM, Sun Z, Sabbagh JJ, Nordhues BA, Koren J, 3rd, et al. (2017) Hsp90 activator Aha1 drives production of pathological tau aggregates. Proc Natl Acad Sci USA 114: 9707–9712

24. Street TO, Lavery LA, Verba KA, Lee C-T, Mayer MP, Agard DA (2012) CrossMonomer Substrate Contacts Reposition the Hsp90 N-Terminal Domain and Prime the Chaperone Activity. J Mol Biol 415: 3–15

25. Weikl T, Muschler P, Richter K, Veit T, Reinstein J, Buchner J (2000) C-terminal regions of Hsp90 are important for trapping the nucleotide during the ATPase cycle. J Mol Biol 303: 583–592

26. Hessling M, Richter K, Buchner J (2009) Dissection of the ATP-induced conformational cycle of the molecular chaperone Hsp90. Nat Struct Mol Biol 16: 287–293

27. Siligardi G, Hu B, Panaretou B, Piper PW, Pearl LH, Prodromou C (2004) Cochaperone Regulation of Conformational Switching in the Hsp90 ATPase Cycle. J Biol Chem 279: 51989–51998

28. Tugarinov V, Kanelis V, Kay LE (2006) Isotope labeling strategies for the study of high-molecular-weight proteins by solution NMR spectroscopy. Nat Protoc 1: 749–754

29. Franke D, Petoukhov MV, Konarev PV, Panjkovich A, Tuukkanen A, Mertens HDT, Kikhney AG, Hajizadeh NR, Franklin JM, Jeffries CM, et al. (2017) ATSAS 2.8: a comprehensive data analysis suite for small-angle scattering from macromolecular solutions. J Appl Crystallogr 50: 1212–1225

## References

1. Oroz J, Chang BJ, Wysoczanski P, Lee CT, Perez-Lara A, Chakraborty P, Hofele RV, Baker JD, Blair LJ, Biernat J, et al. (2018) Structure and pro-toxic mechanism of the human Hsp90/PPIase/Tau complex. Nat Commun 9: 4532

2. Verba K, A., Wang RY-R, Arakawa A, Liu Y, Shirouzu M, Yokoyama S, Agard DA (2016) Atomic structure of Hsp90-Cdc37-Cdk4 reveals that Hsp90 traps and stabilizes an unfolded kinase. Science 352: 1542–1547

3. Retzlaff M, Hagn F, Mitschke L, Hessling M, Gugel F, Kessler H, Richter K, Buchner J (2010) Asymmetric Activation of the Hsp90 Dimer by Its Cochaperone Aha1. Mol Cell 37: 344–354

4. Meyer P, Prodromou C, Liao C, Hu B, Roe SM, Vaughan CK, Vlasic I, Panaretou B, Piper PW, Pearl LH (2004) Structural basis for recruitment of the ATPase activator Aha1 to the Hsp90 chaperone machinery. EMBO J 23: 1402–1410

5. Koulov AK, LaPointe P, Lu B, Razvi A, Coppinger J, Dong M-Q, Matteson J, Laister R, Arrowsmith C, Yates III JR, et al. (2010) Biological and structural basis for Aha1 regultaion of Hsp90 ATPase activity in maintaining proteostasis in the human disease cystic fibrosis. Mol Biol Cell 21: 871–884

